# Impaired Lung BCAA Metabolism Promotes Ferroptosis and Resultant Pulmonary Arterial Hypertension-Associated Hepatopathy

**DOI:** 10.1101/2025.09.03.672819

**Authors:** Madelyn J. Blake, Jason Hong, Adam Brownstein, Christopher J. Rhodes, Jeffrey C. Blake, Ryan A. Moon, Lynn M. Hartweck, Sasha Z. Prisco, Todd Markowski, LeeAnn Higgins, Kevin Murray, Candance Guerrero, Sandra Breuils-Bonnet, Steeve Provencher, Joanna Pepke-Zaba, Luke S Howard, Mark Toshner, Martin R. Wilkins, Sebastien Bonnet, Kurt W. Prins

## Abstract

**Background:** Dysregulated branched chain amino acid (BCAA) homeostasis occurs in pulmonary arterial hypertension (PAH) as BCAA metabolites accumulate and cause metabolic alterations in pulmonary artery smooth muscle cells (PASMC). In other cells, altered BCAA metabolism promotes ferroptosis, a PAH-inducing metabolic pathway. However, the interplay between BCAAs, lung ferroptosis, and PAH is unexplored, as is the impact of PAH severity on liver molecular regulation, a key unknown as recent clinical data highlight the importance of the lung-right heart-liver axis in PAH outcomes.

**Methods:** Human metabolomic and transcriptomic studies examined BCAA metabolism and ferroptosis pathways. The relationship between BCAAs and ferroptotic-phenotypes in PASMCs was evaluated. Multi-omics and physiological analyses evaluated how modulation of BCAA catabolism impacted preclinical PAH multi-organ physiology. Confocal microscopy and proteomic analyses assessed hepatic alterations in human PAH.

**Results:** Metabolomic analyses identified alterations in BCAA metabolites across multiple physiological gradients in patients with pulmonary vascular disease. RNA sequencing demonstrated deficits in the BCAA catabolic and ferroptosis pathways in PAH lungs and smooth muscle cells. *In vitro*, excess BCAAs induced mitochondrial fragmentation, reactive oxygen species generation, and lipid peroxidation in PASMC. Moreover, BCAAs promoted PASMC death, which ferrostatin-1, a ferroptosis antagonist, rescued. BT2, a small-molecule inducer of BCAA catabolism, reduced PAH severity, improved RV function, and enhanced maximal exercise capacity in monocrotaline rats. BT2 blunted pro-ferroptotic changes in lung metabolites and proteins, and combatted peri-vascular complement deposition. In the liver, BT2 blocked mechanical shear stress phenotypes including hepatocyte nuclear expansion and restructured mitochondrial protein regulation and the metabolomic signature. Additionally, a low BCAA diet modestly combatted preclinical PAH severity. Finally, human PAH livers exhibited increased hepatocyte nuclear size and derangements in liver metabolic regulation.

**Conclusions:** Impaired BCAA metabolism promotes PAH via ferroptosis. PAH severity is associated with hepatic pathological shear stress phenotypes and metabolic alterations, which are combatted by a BCAA-targeted therapy.

**Graphical Abstract/Summary Figure:** 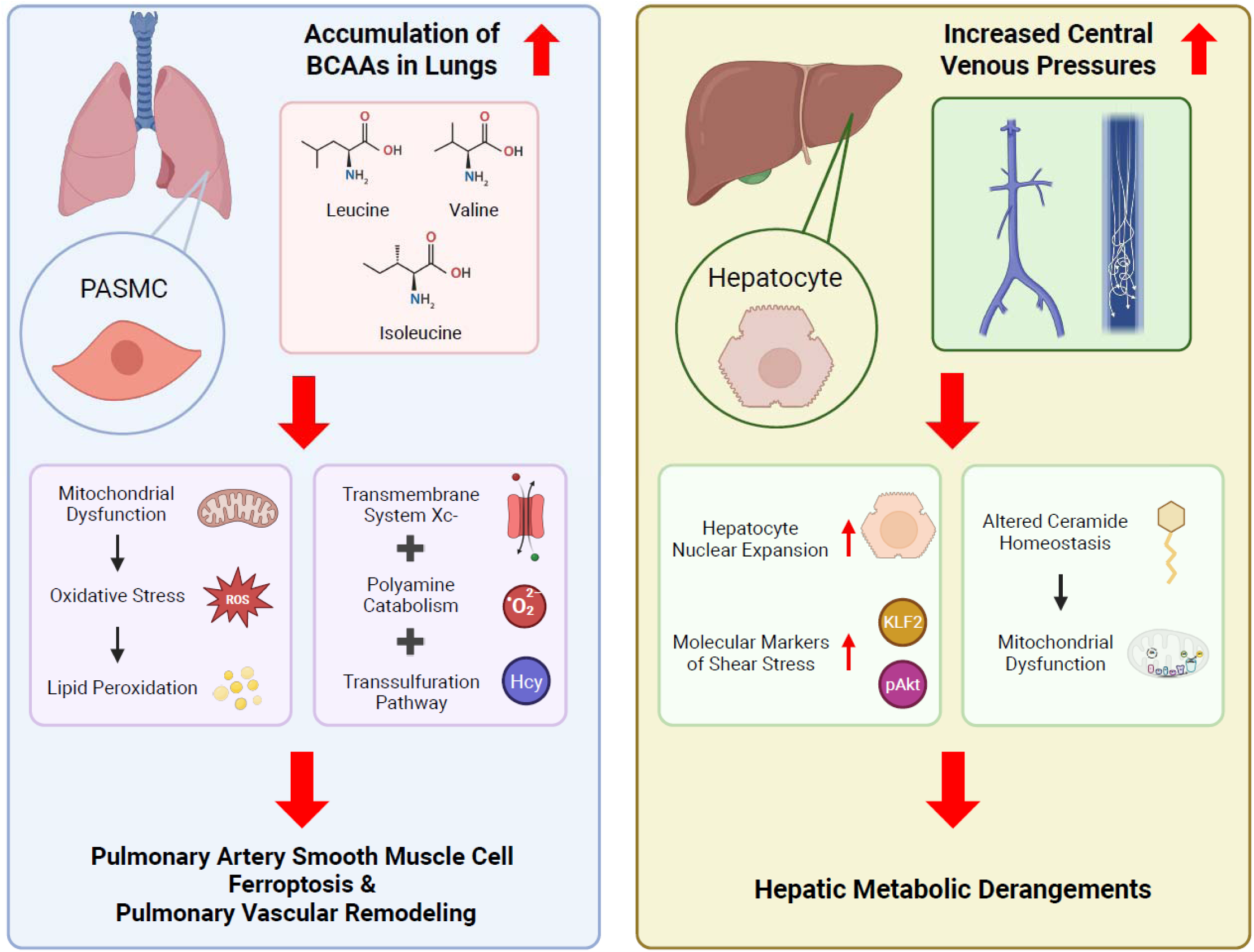

## Introduction

Valine, leucine, and isoleucine are the three branched-chain amino acids (BCAA), and dysregulation of BCAA homeostasis is implicated in multiple cardiovascular diseases^1^. Emerging data suggest impairments in BCAA metabolism contribute to pulmonary arterial hypertension (PAH) via alterations in pulmonary artery smooth muscle cell (PASMC) biology^2^. In particular, the accumulation of branched-chain ketoacids (BCKA), downstream metabolites of BCAAs, aerobically activates hypoxia-inducible factor 1 (HIF1) and induces PAH-like phenotypes in PASMC *in vitro*^2^. However, the potential therapeutic implications of modulating BCAA metabolism in PAH are undefined. Moreover, the relationship between BCAA catabolism and other disrupted metabolic pathways in PAH is unexplored.

Alterations in BCAA catabolism are linked to ferroptosis, a type of cell death triggered by lipid peroxidation due to mitochondrial dysfunction^3^. Inhibition of branched-chain amino acid aminotransferase 2 (BCAT2), an enzyme required for BCAA catabolism, triggers ferroptosis in cancer cells^4^. In cerebral ischemia-perfusion, activation of BCAA oxidation with the small molecule BT2 alleviates neuronal ferroptosis and improves neurological recovery after ischemia-reperfusion injury^4^. In the cisplatin mouse model of chronic kidney disease, upregulation of microRNA 429-3P suppresses BCAA catabolism and promotes renal ferroptosis^5^. Furthermore, BT2-mediated activation of BCAA oxidation reduces multiple measures of ferroptosis in the proximal tubule of cisplatin challenged mice^5^. We^6^ and others^7–9^ have demonstrated that ferroptosis promotes PAH through metabolic and inflammatory mechanisms, but the effects of modulating BCAA homeostasis on PAH severity and associated pulmonary vasculature ferroptosis phenotypes are unknown.

In addition to BCAAs, other amino acids modulate ferroptosis induction. Perhaps the best studied amino acid pathway in ferroptosis is the transmembrane system xc− pathway^10^. Transmembrane system xc− is an amino acid antiporter that exports glutamate and imports cystine, which is then converted to cysteine.

Intracellular cysteine is used to synthesize glutathione, an antioxidant that maintains the anti-ferroptotic protein glutathione peroxidase 4^10^. In addition, the transsulfuration pathway modulates ferroptosis^11^ as it facilitates the conversion of methionine to homocysteine, and then cysteine, which subsequently feeds back to regulate glutathione peroxidase 4^10^. Moreover, recent data show arginine has a pro-ferroptotic function through its downstream metabolites, ornithine and polyamines^12^. At present, the interplay between BCAAs, system xc−, transsulfuration, and arginine metabolites in PAH lungs is not well defined.

Disrupted hepatic function is a risk factor for mortality in PAH^13^ as slight alterations in hepatic function tests are highly predictive of survival^14^. Furthermore, heightened liver stiffness is observed in PAH, and liver stiffness correlates with PAH severity^15^. Currently, the mechanisms underlying hepatic dysfunction in PAH are poorly understood. Disrupted and turbulent venous flow due to right heart failure is a likely contributor^13^, however, a detailed evaluation of altered cellular mechanical stress phenotypes due to disturbed venous flow in PAH is lacking. Two classic and repeatedly identified cellular phenotypes of cells exposed to excess mechanical stress/shear stress are expansion in nuclear size^16,17^ and disrupted mitochondrial regulation^16^.

Thus, these two phenotypes may serve as readouts for mechanical stress, however the impact of PAH on liver nuclear morphology and mitochondrial and metabolic regulation are not well described.

Here, we addressed the interplay between BCAA catabolism, PASMC ferroptosis, amino acid homeostasis, PAH severity, and hepatic molecular phenotypes by evaluating BCAA metabolite levels in humans with pulmonary vascular disease, probing BCAA and ferroptosis pathway dysregulation in human lungs and PASMC using bulk and single nucleus RNAseq, and defining how excess BCAAs modulated ferroptosis phenotypes in human PASMC *in vitro*. Then, we determined how small molecule-mediated augmentation of BCAA catabolism and a low BCAA diet impacted PAH severity, lung ferroptosis phenotypes, and liver function in the monocrotaline rat model of PAH. Finally, we examined how PAH impacted liver histopathology and proteomic regulation in human liver samples.

## Methods

### Data Availability Statement

Data sets, analyses, and study materials will be made available to other researchers upon request. RNAseq data will be made publicly available upon publication of Brownstein *et. al.* (manuscript in preparation). All data for omics experiments are publicly available. The mass spectrometry (MS) proteomics data and metabolomics/lipidomics are available at Zenodo: doi:10.5281/zenodo.15832197

### Human Liver Tissue

Human liver tissue samples were collected at Laval University. Tissues were obtained from patients who had previously given signed informed consent. Control and PAH tissues were collected from warm autopsies. Prior to tissue collection, PAH diagnosis was confirmed by right heart catheterization. All tissues were stored at −80°C prior to analysis.

### Rodent Studies

Small-molecule mediated induction of BCAA catabolism in the monocrotaline (MCT, Sigma-Aldrich) rat PAH model was evaluated in male Sprague-Dawley (Charles River) rats. Rodents at 7-8 weeks of age were randomly assigned to one of three groups: 1. Control rats given a one-time subcutaneous injection of phosphate buffered saline (*n*=10), 2. Rats injected subcutaneously with 60 mg/kg MCT and then treated with daily intraperitoneal injections of vehicle (2% dimethyl sulfoxide, 50% polyethylene glycol, 5% Tween 80, and 43% double distilled water) starting two weeks after MCT injection (*n*=20), and 3. MCT rats treated with daily intraperitoneal BT2 (20 mg/kg, MedChemExpress) starting two weeks after MCT injection (*n*=15). A low-BCAA dietary intervention was also evaluated in the MCT PAH model using male Sprague-Dawley rats. Rodents at 7-8 weeks of age were randomly assigned to one of two groups: 1. Rats subcutaneously injected with 60 mg/kg MCT and then fed a standard chow diet (Teklad, 2918) (*n*=10) or 2. MCT rats fed a standard chow diet (Teklad, 2918) for 14 days, before switching to a low BCAA chow (Envigo, TD.140714) for 10 days (*n*=10). End-point analysis was performed 25 days post MCT injection with primary physiological read-out being right ventricular systolic pressure (RVSP). Females were not evaluated due to the more robust phenotype observed in males. Sample size was estimated at *n*=20 based on previous experience with phenotypic variation, rodent death due to MCT, and statistical power^18^. Three MCT-Vehicle rodents and one MCT-standard diet rodent in the dietary experiment died before end-point analysis. All rodent studies were approved by the University of Minnesota Institutional Animal Care and Use Committee.

### Maximal Exercise Testing on Rodent Treadmill

Testing was performed on a four-lane modular rodent treadmill (AccuPacer, Omnitech Electronics Inc.) with individual metabolic chambers and a built-in gas analyzer system that recorded metabolic profiles (VO_2_ max, respiratory exchange rate, etc.) real-time. Rodents were habituated to the treadmill one week prior to end-point analysis using an adapted protocol from Zhang et. al.^19^ with three episodes of gradual running building up to a final speed of 15m/min at a 15-degree incline for 10 minutes. End-point maximal exercise testing used the exercise testing protocol previously described by Bedford et. al^20^.

### Cardiovascular Phenotyping

Echocardiography and closed-chest hemodynamic analyses quantified RV function and pulmonary vascular disease as we have previously described^18.21^.

### PASMC BCAA Treatment

Human PASMC (ATCC PCS-100-023) were grown with Sigma Basic Eagle Medium (Sigma, B1522-500) with supplements (Lonza, CC-3182) and passaged with subculture reagents (Lonza CC-5034). PASMC cells were received at passage 2 (P2) and used at P4-6. BCAA media was produced by supplementing Sigma Basic Eagle Medium with 0.6 mM isoleucine (Sigma Aldrich, I2752), 0.6 mM leucine (Sigma Aldrich, L8000), and 0.6 mM valine (Sigma Aldrich, V0500), such that each BCAA had a final concentration of 0.8 mM.

### PASMC Mitochondrial Morphology Analysis

After incubation with BCAA media for 72 hours, PASMC were stained with MitoTracker Orange (Thermo Scientific, M7514), fixed in 4% paraformaldehyde, and then mounted in Prolong Mountant with NucBlue (Thermo Scientific). Images were collected on a Zeiss LSM 900 Airyscan 2.0 microscope and then blindly analyzed by JCB using the Mitochondrial Network Analysis Plug-In59 (https://github.com/StuartLab) in FIJI.

### PASMC Live-Cell Experiments

70-80% confluent PASMC were cultured in BCAA media for 72 hours before being subjected to live-cell imaging on Zeiss LSM 900 Airyscan 2.0 microscope. For assessment of mitoROS production, BCAA-treated PASMC were treated with 500nM MitoSox (ThermoFisher). For mitochondrial membrane potential staining, BCAA-treated PASMC were treated with 200nM TMRE (Abcam, ab113852). For lipid peroxidation staining, PASMC were co-treated with 50uM oleate-BSA (Cayman) and 50uM palmitate-BSA (Cayman) during the BCAA treatment period. The BCAA ferrostatin-1 treatment PASMC received 5μM ferrostatin-1 for 24 hours prior to staining. Following lipid and BCAA treatment, PASMC were stained with 0.5µM BODIPY (ThermoFisher). All stains were incubated with PASMCs for 30 minutes at 37C before a gentle washout with media and all live-cell imaging was completed within 4 hours of the washout step to ensure optimal results. MitoSox, TRME, and BODIPY analyses were blindly analyzed by JCB in FIJI.

### PASMC Cell Death Assay

Susceptibility to cell death in BCAA-treated PASMC was probed by treating with BCAAs for 72 hours. The BCAA ferrostatin-1 PASMCs received 5μM ferrostatin-1 for the last 24 hours. Following treatment, the PASMC were stained with 200uL of 0.4% Trypan Blue for 10 minutes, before the stain was gently washed out. Stained PASMC were then fixed in 4% paraformaldehyde and mounted in Prolong Mountant with NucBlue (Thermo Scientific). Images were collected on a Zeiss LSM 900 Airyscan 2.0 microscope and then blindly analyzed by MJB.

### Lung and Liver Mitochondrial Proteomics Analysis

Mitochondrial enrichments (Abcam) from lung and liver tissue were performed according to manufactures instructions and were subjected to TMT16-plex (ThermoFisher) labeling and quantitative proteomics using Proteome Discover Software as previously described^18,21^. The lung mitochondrial-enriched proteome in *n*=5 control, *n*=5 MCT-Vehicle, and *n*=6 MCT-BT2 animals were analyzed. For the rodent liver specimens, *n*=5 control, *n*=5 MCT-Vehicle, and *n*=6 MCT-BT2 animals were included. Human liver samples consisted of *n*=4 control and *n*=3 PAH patients.

### Lung and Liver Metabolomics/Lipidomics

Global metabolomics analysis of frozen lung specimens (*n*=5 Control, *n*=13 MCT-Vehicle, and *n*=6 MCT-BT2) and frozen liver specimens (*n*=8 Control, *n*=8 MCT-Vehicle, and *n*=8 MCT-BT2) was performed at the University of Minnesota Center for Metabolomics and Proteomics using the Biocrates’ MxP® Quant 500 kit. Samples were homogenized in 2.0 mL Precelly standard tubes.

Samples were centrifuged at 10,000*g* for 5 minutes at 4°C and supernatant was collected. 10µL of the extract was loaded and classes of metabolites were determined using 50µL of 1:1:1:0.16 water:EtOH:pyridine:phenyl isothiocyanate solution and incubated for an hour. ABSciex QTRAP 5500 triple-quadrupole (Farmington, MA, USA) mass spectrometer was used to perform the metabolomic assays.

### Serum Metabolomics/Lipidomics

Global metabolomics analysis of frozen serum specimens from the BT2 study (*n*=10 control, *n*=17 MCT-Vehicle, and *n*=10 MCT-BT2) and the dietary intervention study (*n*=9 MCT-standard and *n*=9 MCT-low BCAA) was performed at the University of Minnesota Center for Metabolomics and Proteomics using the Biocrates’ MxP® Quant 500 kit. 10µL of the thawed serum was loaded and classes of metabolites were determined using 50µL of 1:1:1:0.16 water:EtOH:pyridine:phenyl isothiocyanate solution and incubated for an hour. ABSciex QTRAP 5500 triple-quadrupole (Farmington, MA, USA) mass spectrometer was used to perform the metabolomic assays.

### Rodent Immunohistochemistry

Rat lung (C3 (Abcam, ab200999): α_sm_-actin (Sigma Aldrich, 1A4): transferrin receptor (Abcam, ab214039), 1:50 dilution overnight) sections were subjected to antigen retrieval (Reveal Decloaker, BioCare), incubated with primary antibody and processed for immunofluorescence analysis. Rat and human liver sections were stained with wheat germ agglutinin (WGA) and 4′,6-diamidino-2-phenylindole (DAPI) to delineate hepatocyte nuclear morphology. Confocal images were collected on a Zeiss LSM 900 Airyscan 2.0 and blindly analyzed by JCB with FIJI (NIH).

### Western Blots

Protein was isolated from flash-frozen lung (*n*=5 control, *n*=5 MCT-Vehicle, and *n*=5 MCT-BT2) and liver (*n*=4 control, *n*=4 MCT-Vehicle, and *n*=4 MCT-BT2) in sodium dodecylsulfate (SDS) buffer. 25 μg of protein extracts were used for immunoblotting and blots were imaged using the Odyssey Infrared Imaging system as previously described^21,22^. Post-transfer SDS-PAGE gels were stained with Coomassie brilliant blue (CBB). Rat lung protein extracts were exposed to the following primary antibodies: mTOR (Cell Signaling, mAb 2983), phospho-mTOR (Cell Signaling, mAb 2971), p70S6K (Cell Signaling, mAb 14130), and phospho-p70S6K (Cell Signaling, mAb 9204) at a 1:250 dilution overnight. Primary antibodies used for rat liver extracts were: KLF2 (Abcam, ab314430), Akt (Cell Signaling, mAb 9272), and phospho-Akt (Cell Signaling, mAb 9271) at a 1:250 dilution overnight. For all blots, the secondary antibody used was 800CW Anti-Rabbit IgG Goat Secondary Antibody (LiCORbio, 926-3221) at a 1:5000 dilution for 1 hour.

### UK PAH Patient Plasma Metabolite Gradient Methods

Patients were recruited from Hammersmith and Royal Papworth Hospital PH Services with written informed consent and local ethical approvals. Plasma samples were collected during right heart catheterization from the radial artery (ART), superior vena cava (SVC), and pulmonary artery (PA) and snap frozen in liquid nitrogen and stored at −80°C until analysis of metabolites by LC-MS/MS by Metabolon. Normalized and z-scored BCAA metabolite levels were compared between bleed sites by paired t-tests.

### UCLA Data/Analysis

Sample-level enrichment scores of BCAA catabolism and ferroptosis pathways were determined using the GSVA R package^23^ (v1.44) in a bulk lung RNA-seq cohort of 96 PAH and 52 control human lungs from the Pulmonary Hypertension Breakthrough Initiative^24^ (PHBI; GSE254617) with KEGG-derived gene sets for BCAA catabolism and ferroptosis. As previously described^24^, SMC abundance was deconvoluted from bulk lung RNA-seq samples using CIBERSORTx^25^ and an integrated human lung single-cell reference atlas. A targeted analysis of SMCs was also performed using a subset of a larger single-nucleus RNA-seq study of 42 PAH and 25 control human lungs from PHBI (Brownstein et al., manuscript in preparation; see also Hong *et al*^26^., *Circulation* 2024;150:Suppl_1:A4137829). SMCs were identified by canonical markers, and pathway activity scores for BCAA catabolism and ferroptosis were calculated per cell using Seurat’s AddModuleScore function^27^ with KEGG-derived gene sets. Scores were averaged per patient prior to comparison between PAH and control groups.

### Statistical Analysis

Statistical analyses were performed using GraphPad Prism 10. Normality of data was determined using the Shapiro-Wilk test. If data were normally distributed and there was equal variance as determined using the Brown-Forsythe test, one-way analysis of variance with Tukey’s multiple-comparisons test was performed. If there was unequal variance, Brown-Forsythe and Welch analysis of variance with Dunnett multiple-comparisons test was completed. If the data were not normally distributed, the Kruskal-Wallis test and Dunn’s multiple-comparisons test were used when comparing two groups and the Mann Whitney U test for comparing two groups. Graphs display the mean or median value and all individual values. *p*-values of <0.05 were considered to indicate statistical significance. Exact *p*-values are provided, except in the case of *p*-values < 1.0e^-15^, after which Prism 10 does not provide exact values. Hierarchical cluster analyses, sparse least discriminant analysis, random forest classification, and correlational heatmapping were performed using MetaboAnalyst (https://www.metaboanalyst.ca/). The relative abundance of each protein was determined using Proteome Discover. Transcripts and proteins that were significantly correlated (*p*<0.05) with central venous pressures and right ventricular systolic pressure were processed for Kyoto Encyclopedia of Genes and Genomes (KEGG) and WikiPathways (Wiki) using ShinyGO 0.76.3 (http://bioinformatics.sdstate.edu/go/).

## Results

### Metabolomics Analysis Identified Altered BCAA Metabolites in Human Pulmonary Vascular Disease

To determine if BCAA homeostasis was disrupted in patients with pulmonary vascular disease, we probed BCAAs and BCAA catabolite levels (**Figure 1A**) in arterial (ART), superior vena cava (SVC), and pulmonary artery (PA) blood samples and gradients across those sites from patients with PAH (*n*=18) or chronic thromboembolic pulmonary hypertension (*n*=68) (**Figure 1B**). We observed alterations in BCAAs as the isoleucine SVC-PA gradient and leucine PA-ART gradient were significantly increased in pulmonary hypertension patients (**Figure 1C**). In addition, further downstream metabolites in the BCAA pathway were modified as the SVC-PA gradients of 2-hydyroxy-3-methylvalerate, 2-methylbutyrylcarnitine, and 3-hydroxy-2-ethylpropionate were altered in patients with pulmonary vascular disease (**Figure 1C)**. Finally, the PA-ART gradients of 3-hydroxy-2-ethylpropionate and ethylmalonate were deranged in the diseased patients (**Figure 1C**).

**Figure 1:**
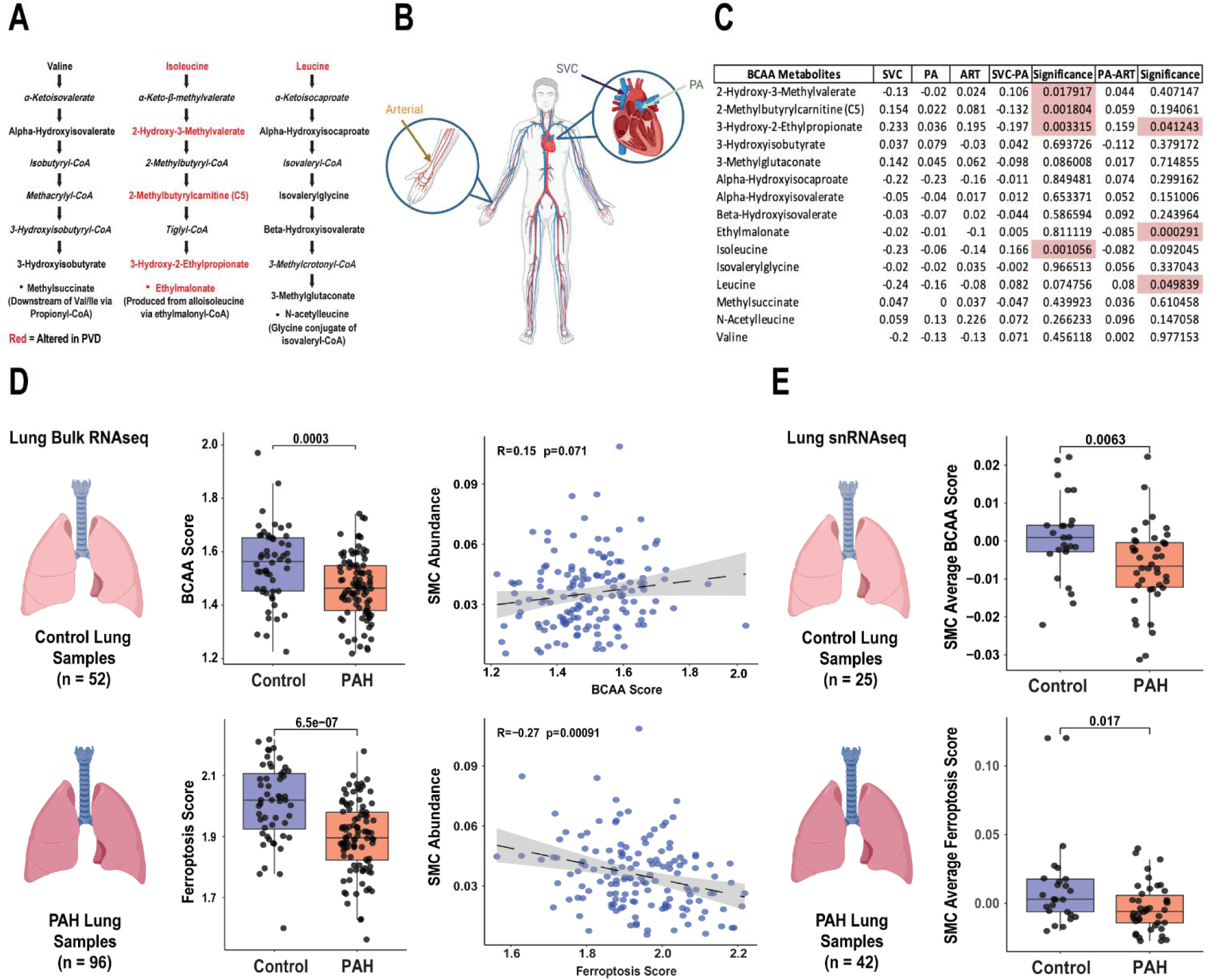
Human metabolomics, bulk RNAseq, and snRNAseq data linked altered BCAA catabolism and ferroptosis to pulmonary vascular disease. (A) Schematic representation of BCAA metabolism with associated metabolites. (B) Graphical depiction of gradient blood sampling for human metabolomics analysis. (C) Summary of BCAA-associated metabolite alterations at each individual site and gradients with associated p-values in patients with pulmonary vascular disease. (D) Lung bulk RNAseq demonstrating alterations in BCAA catabolism and ferroptosis pathways in PAH lungs, with SMC deconvolution displaying a positive, but statistically insignificant, trend between SMC abundance and BCAA catabolism, and a significant negative correlation between SMC abundance and ferroptosis activity. p-values determined by two-sided Wilcoxon rank-sum test for group comparisons and Pearson correlation for associations. (E) Lung snRNAseq mirrors bulk RNAseq with PAH SMCs showing alterations in both BCAA catabolism and ferroptosis activity scores relative to controls. p-values determined by two-sided Wilcoxon rank-sum test.

### Bulk and snRNAseq Data from Human PAH Samples Demonstrated Dysregulation of the BCAA Catabolic Pathway and Ferroptosis in Human SMC

Then, we evaluated the BCAA and ferroptosis pathways in a lung bulk RNAseq analysis comparing 52 control samples to 96 PAH patients. There were significant reductions in both the BCAA catabolism and ferroptosis pathway scores in PAH lungs as compared to controls (**Figure 1D**). Then, we probed the relationship between BCAA score and deconvoluted smooth muscle cell (SMC) relative abundance and found a nonsignificant (*p*=0.071) association between SMC abundance and BCAA score (**Figure 1D**). However, there was a highly significant and negative association between ferroptosis score and SMC abundance (*p*=0.0009) (**Figure 1D**). A targeted analysis of BCAA and ferroptosis pathways in SMCs was performed in human lung snRNAseq data from 25 control and 42 PAH patients (Brownstein *et. al.*, manuscript in preparation). In agreement with the bulk RNAseq data, both the BCAA and ferroptosis scores were significantly lower in PAH SMC than controls (**Figure 1E**). Thus, these human data supported the hypothesis that human PAH was marked by alterations in BCAA and ferroptosis regulation in SMC.

### Excess BCAAs Promoted a Pro-Ferroptotic Phenotype in Human PASMC In Vitro

To directly examine the link between BCAAs and ferroptosis in human PASMC, we evaluated how excess BCAAs modulated PASMC ferroptotic phenotypes *in vitro*. Heightened BCAAs induced mitochondrial fragmentation (**Figure 2A**), altered mitochondrial membrane potential (**Figure 2B**), and triggered mitochondrial reactive oxygen species generation (**Figure 2C**). Moreover, elevated BCAAs increased lipid peroxidation levels, the trigger for ferroptosis^28^ (**Figure 2D**). Finally, PASMCs treated with BCAAs exhibited heightened sensitivity to cell death, which was rescued by ferrostatin-1, a small molecule inhibitor of ferroptosis^29^ (**Figure 2E**), suggesting that excess BCAAs promoted a pro-ferroptotic phenotype in human PASMC.

**Figure 2:**
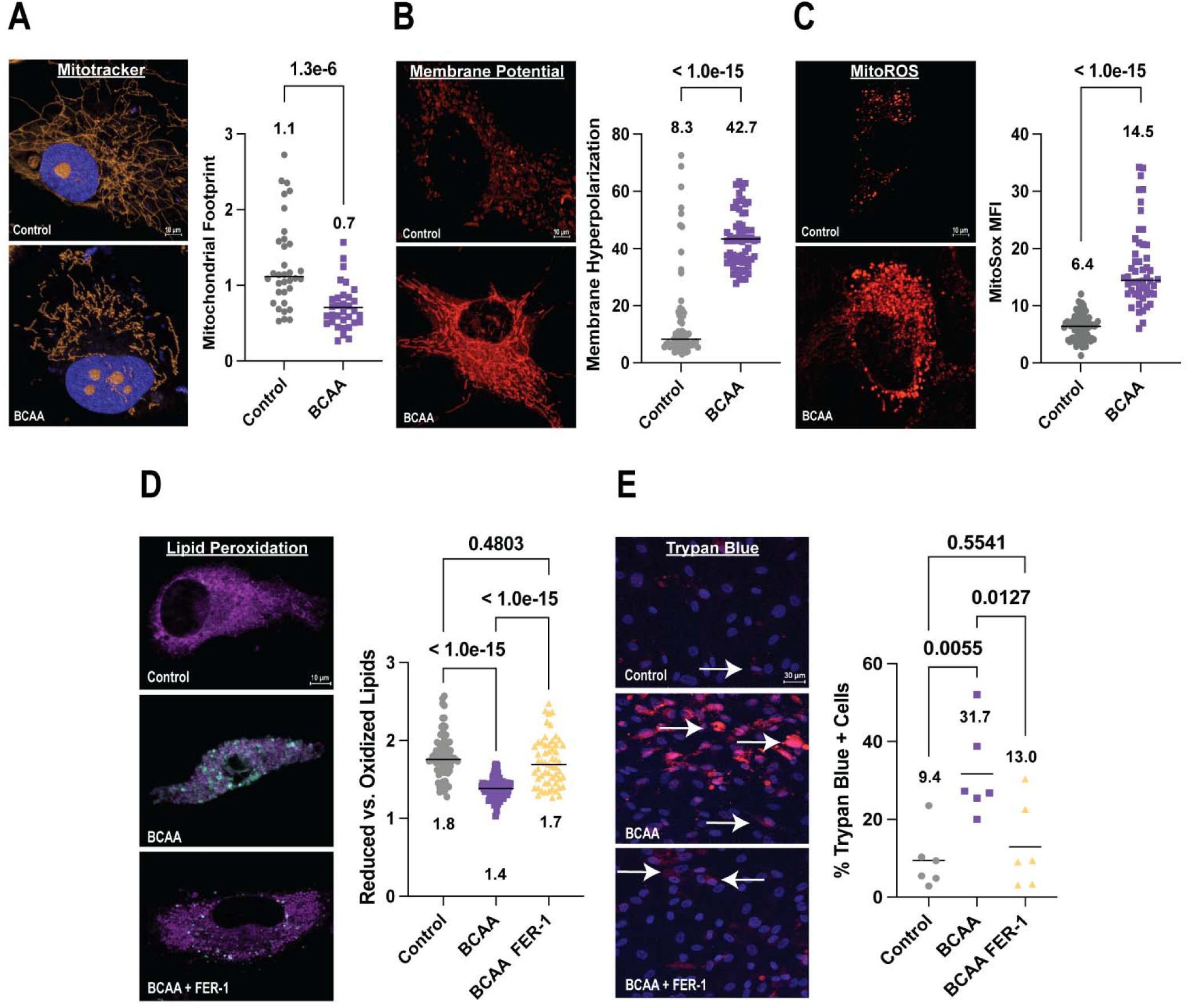
**Excess BCAAs promoted a pro-ferroptotic phenotype in human PASMC**. (A) Representative confocal micrographs of PASMC stained with MitoTracker Orange and quantification of mitochondrial organization in control and BCAA-treated PASMC (Control PASMC [Con]: 1.1±0.6, BCAA-Treated PASMC [BCAA]: 0.7±0.3); *p*-values determined by Mann-Whitney U-test. (B) Confocal micrographs of TRME-stained PASMC in control and BCAA-treated media, with corresponding quantification of mitochondrial membrane hyperpolarization, indicated by fluorescence intensity (Con: 8.3±17.2, BCAA: 42.7±9.7). *p*-values determined by Mann-Whitney U-test. (C) Excess BCAAs increase mitochondrial ROS, demonstrated by confocal micrographs of control and BCAA-treated PASMCs stained with MitoSox Red (Con: 6.4±2.1, BCAA: 14.5±6.9 MFI). *p*-values determined by Mann-Whitney U-test. (D) Representative confocal micrographs of control, BCAA-treated, and BCAA and ferrostatin-1-treated PASMCs incubated with 50 μM oleate and 50 μM palmitate and stained for lipid peroxidation using BODIPY (Con: 1.8±0.3, BCAA: 1.4±0.1, BCAA-treated with 5 μM ferrostatin-1 [BCAA-FER1]: 1.7±0.3). *p*-values determined by Kruskal-Wallis test and Dunn’s multiple comparisons test. (E) Incubation with BCAAs induces ferroptotic cell death, demonstrated by viability staining of control, BCAA-treated, and BCAA-treated with ferrostatin-1 PASMCs via Trypan Blue (Con: 9.4±7.5, BCAA: 31.7±11.7, BCAA-FER1: 13.0±11.1). White arrows indicate trypan blue-positive cells. *p*-values determined by ordinary one-way *ANOVA* with Tukey’s multiple comparison test.

### Activation of BCAA Catabolism with BT2 Mitigated PAH Severity, Improved RV Function, and Enhanced Exercise Capacity in Monocrotaline Rats

To understand how modulating BCAA metabolism impacted PAH pathobiology, we treated MCT rats with the small molecule BT2, a well-validated activator of BCAA catabolism^30^. Hemodynamic evaluation demonstrated BT2 counteracted PAH as right ventricular systolic pressure (RVSP), mean pulmonary arterial pressure (mPAP), and total pulmonary resistance index (TPRi) all decreased compared to MCT-Vehicle rodents (**Figure 3 A-C**). Histological analysis confirmed BT2 treatment suppressed small pulmonary arterial remodeling (**Figure 3D**). There was blunted right ventricular (RV) hypertrophy, both at the organ-level demonstrated by a decreased Fulton index (**Figure 3E**) and at the cardiomyocyte-level with smaller cardiomyocyte cross-sectional area (**Figure 3F**). In addition, BT2 treatment reduced RV fibrosis (**Figure 3G**) and augmented right ventricular function as tricuspid annular plane systolic excursion (TAPSE), RV free wall thickening, and cardiac output normalized to body weight were all significantly improved (**Figure 3 H-J**).

**Figure 3:**
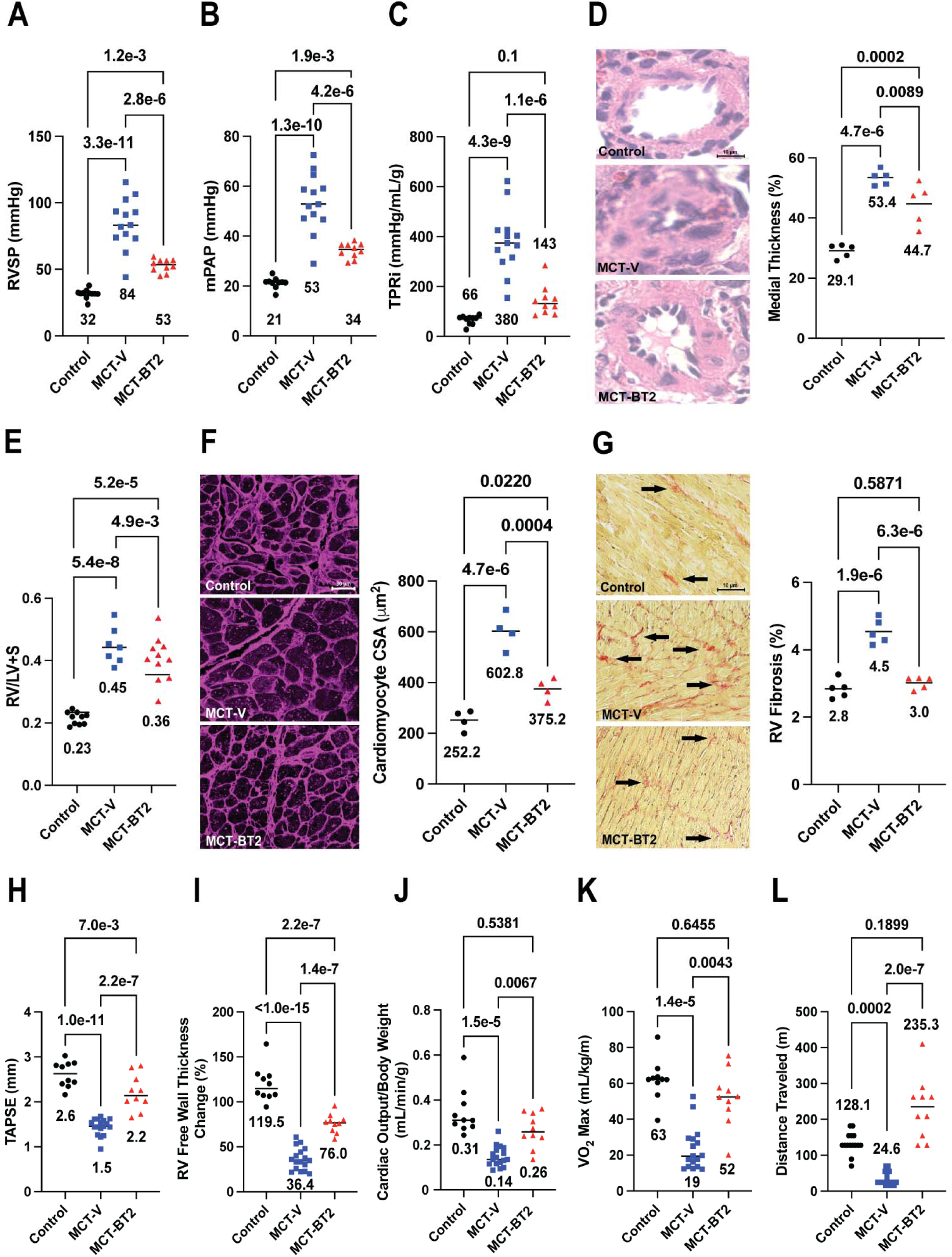
**BT2-mediated activation of BCAA catabolism mitigated cardiopulmonary alterations in monocrotaline rats**. (A) BT2 (20mg/kg) treatment reduced right ventricular systolic pressure (RVSP, control [Con]: 32±3 [n=10], MCT-Vehicle [MCT-V]: 84±19 [n=13], MCT-BT2: 53±5 [n=10] mmHg), (B) mean pulmonary artery pressure (mPAP, Con: 21±2, MCT-V: 53±12, MCT-BT2: 34±3 mmHg), and (C) total pulmonary resistance index (TPRi, Con: 66±18, MCT-V: 380±126, MCT-BT2: 143±60 mmHg/mL/g); *p*-values determined by one-way *ANOVA* with Tukey’s multiple comparison test. (D) Small vessel pulmonary arterial remodeling was mitigated by BT2 (median values averaged from 5 animals per group: Con: 29.1±2.3, MCT-V: 53.4±2.5, MCT-BT2: 44.7±6.9 %; *p*-values determined by ordinary one-way *ANOVA* with Tukey’s multiple comparison test). Lung tissue was fixed in 10% formalin, paraffin embedded, sectioned at 4-µm, and stained with hematoxylin and eosin. (E) BT2 diminished RV hypertrophy at the organ level as assessed by the Fulton index (Con: 0.23±0.02, MCT-V: 0.45±0.06, MCT-BT2: 0.36±0.07; *p*-values determined by ordinary one-way *ANOVA* with Tukey’s multiple comparison test) and (F) at the cardiomyocyte level (median cross sectional area values averaged from 5 animals per group: Con: 252.2±39.3, MCT-V: 602.8±70.3, MCT-BT2: 375.2±41.8 μm^2^). RV sections were stained with hematoxylin and eosin. (G) RV fibrosis was also mitigated by BT2 treatment (Con: 2.8±0.3, MCT-V: 4.5±0.4, MCT-BT2: 3.0±0.2 %; *p*-values determined by ordinary one-way *ANOVA* with Tukey’s multiple comparison test). RV sections were stained with Picrosirius Red. Arrows indicate areas of fibrosis. Inducing BCAA catabolism augmented (H) tricuspid annual plane systolic excursion (TAPSE, Con: 2.6±0.3, MCT-V: 1.5±0.2, MCT-BT2: 2.2±0.4 mm) and (I) right ventricular free wall thickening (Con: 119.5±19.5, MCT-V: 36.4±12.5, MCT-BT2: 76.0±10.4%); *p*-values determined by ordinary one-way *ANOVA* with Tukey’s multiple comparison test. (J) BT2 treatment also improved cardiac output normalized to body weight (Con: 0.31±0.10, MCT-V: 0.14±0.04, MCT-BT2: 0.26±0.1 mL/min/g; *p*-values determined by Kruskal Wallis test with Dunn’s multiple comparison test). (K) End-point treadmill assessments demonstrated BT2 improved maximal exercise capacity (VO_2_ max, Con: 63±12, MCT-V: 19±12, MCT-BT2: 52±16 mL/kg/m), and total distance traveled on the treadmill (Con: 128.1±29.5, MCT-V: 24.6±19.1, MCT-BT2: 235.3±85.8 m); *p*-values determined Kruskal Wallis test with Dunn’s multiple comparison test.

Maximal right atrial pressures were also minimized in MCT-BT2 animals (Con: 6.0±2.0, MCT-Vehicle: 14.0±3.0, MCT-BT2: 8.0±3.0 mmHg), suggesting an enhancement of RV diastolic function. Finally, we showed BT2 impacted systemic biology as it augmented maximal oxygen consumption rates and total distance ran on treadmill in rodents subjected to maximal exercise testing (**Figure 3K and L**).

### BT2 Treatment Reduced Lung BCAA Levels and Restructured BCAA Metabolic Proteins

Next, we evaluated how BT2 modulated lung BCAA levels and BCAA catabolizing proteins using a combination of metabolomics and proteomics analyses. First, analysis of proteins that were significantly altered in the lungs of the three groups identified BCAA catabolism, further suggesting a link between BCAA, metabolic pathways, and PAH pathobiology (**Supplemental Figure 1A**). When we specifically profiled BCAA metabolism, we found BT2 treatment induced an intermediate phenotype between control and MCT-Vehicle lungs as it mitigated downregulation of several BCAA catabolic proteins (**Supplemental Figure 1B**). Then, metabolomics data demonstrated BT2 significantly reduced lung levels of all three BCAAs (**Supplemental Figure 1C**). Notably, MCT-BT2 animals also had a parallel decrease in serum BCAAs (**Supplemental Figure 1D**). Finally, we evaluated another readout of BCAA accumulation, the mTORC1 pathway in lung samples from the three experimental groups. BT2 blunted mTORC1 induction as lung levels of phosphorylated mTOR and phosphorylated S6 kinase were reduced by BT2 (**Supplemental Figure 1E**).

### BT2 Treatment Suppressed Multiple Pro-Ferroptotic Phenotypes in the Lungs of Monocrotaline Rats

To better examine the link between altered BCAA catabolism and ferroptosis, we evaluated both lung metabolomics and proteomics data. Random forest classification identified arginine, ornithine, sulfur-containing amino acids, calculated rate of homocysteine synthesis (homocysteine/methionine), and homocysteine as important variables that differentiated the three groups (**Figure 4A**). Arginine and ornithine are members of the polyamine catabolic pathway^31^, while cysteine, methionine, and homocysteine are related to the transmembrane system xc^−^ and transsulfuration pathways^32,33^, all of which are implicated in ferroptosis^12,34,35^ (**Figure 4B**). MCT lungs had elevated levels of arginine, ornithine, methionine, cysteine, homocysteine, which BT2 diminished (**Figure 4C**). Importantly, the observed increases in arginine, cysteine, and homocysteine were lung enriched as these metabolites were not significantly altered in the serum. (**Supplemental Figure 2A**).

**Figure 4:**
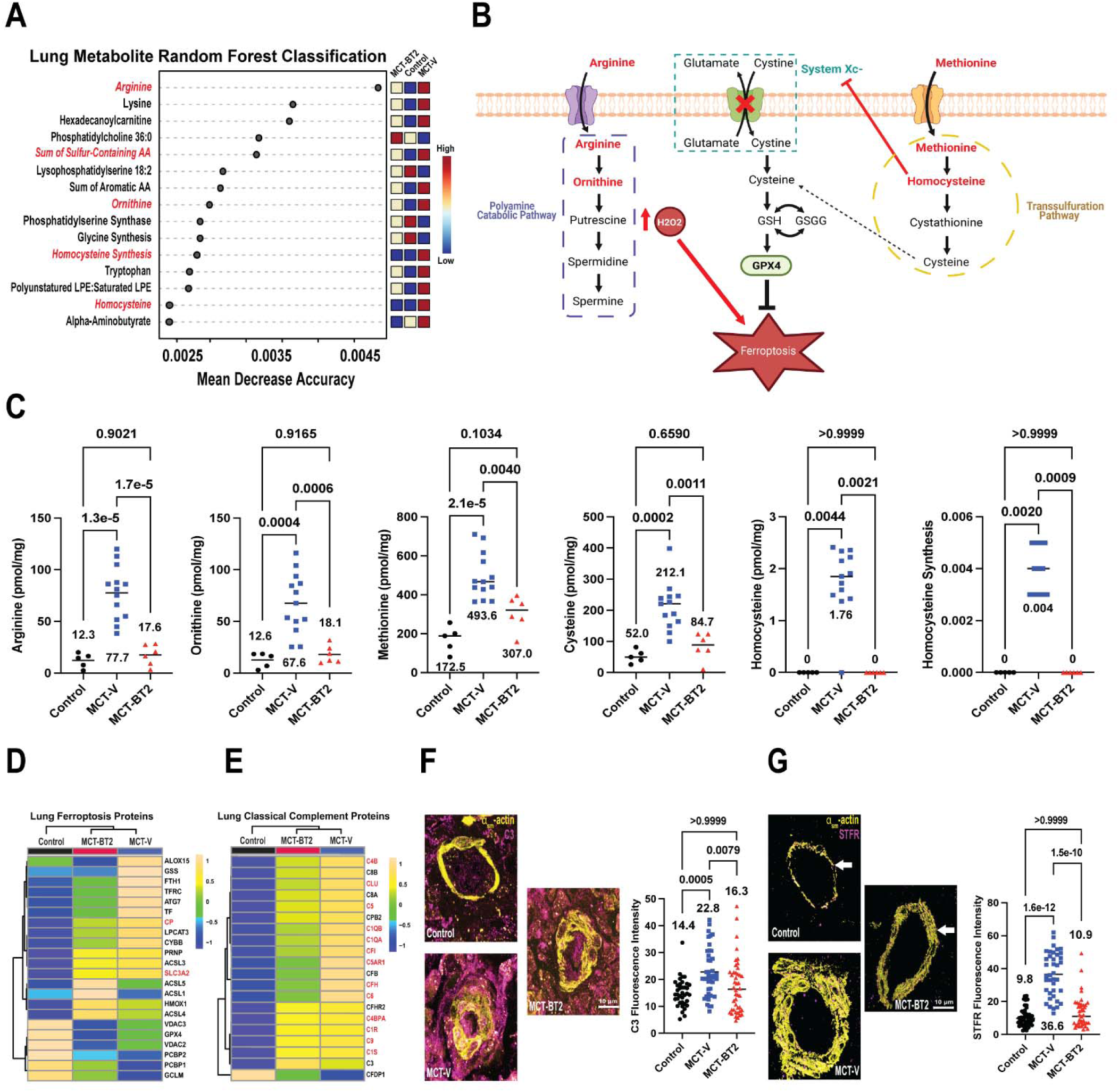
**BT2 combatted a pro-ferroptotic molecular signature in the lungs of monocrotaline rats**. (A) Random forest classification of top lung metabolites and lipids distinguishing control, MCT-V, and MCT-BT2. (B) Schematic representation of amino acid-induced ferroptosis, consisting of the polyamine catabolic pathway, transmembrane system xc^−^, and the transsulfuration pathway. (C) BT2 treatment significantly reduced lung concentrations of arginine (Con: 12.3±7.1, MCT-V: 77.7±25.8, MCT-BT2: 17.6±10.0 pmol/mg), ornithine (Con: 12.6±7.3, MCT-V: 67.6±29.0, MCT-BT2: 18.1±8.6 pmol/mg), methionine (Con: 172.5±67.2, MCT-V: 493.6±118.2, MCT-BT2: 307.0±87.0 pmol/mg), cysteine (Con: 52.0±21.3, MCT-V: 212.1±75.3, MCT-BT2: 84.7±43.3 pmol/mg), homocysteine (Con: 0±0, MCT-V: 1.76±0.64, MCT-BT2: 0±0 pmol/mg), and the calculated rate of homocysteine synthesis (Con: 0±0, MCT-V: 0.004±0.0009, MCT-BT2: 0±0). *p*-values for arginine, ornithine, methionine, and cysteine determined by ordinary one-way *ANOVA* with Tukey’s multiple comparison test, Kruskal-Wallis test and Dunn’s multiple comparison test for homocysteine and homocysteine synthesis. Hierarchical cluster analyses of proteins in the (D) ferroptosis and (E) classical complement pathways, red text denotes statistically significant proteins with *p*-values determined by ordinary one-way *ANOVA* with Tukey’s multiple comparison test. Representative confocal micrographs show increased (F) complement deposition (Con: 14.4±5.2, MCT-V: 22.8±9.0, MCT-BT2: 16.3±9.9 C3 fluorescence intensity) and (G) soluble transferrin receptor concentration (Con: 9.8±5.6, MCT-V: 36.6±13.5, MCT-BT2: 10.9±9.7 STFR fluorescence intensity) around the pulmonary vasculature in MCT-V animals, which was mitigated by BT2 treatment. *p*-values determined by the Kruskal-Wallis test and Dunn’s multiple comparison test.

When probing lung proteomics data, we found multiple ferroptosis proteins were upregulated in MCT-Vehicle rats, which BT2 combatted (**Figure 4D**). Next, we further leveraged our proteomics data to evaluate complement regulation as complement deposition is a marker of lung ferroptosis^6^. Proteomic analysis revealed BT2 partially negated the accumulation of several complement proteins in the lungs (**Figure 4E**). Likewise, immunohistochemistry analysis revealed perivascular complement deposition was present in MCT lungs but reduced with BT2 (**Figure 4F**). Finally, PASMC-specific expression of transferrin receptor, a validated histological marker of ferroptotsis^36^, was increased in MCT rats but reduced by BT2 treatment (**Figure 4G**). The summation of our proteomics, metabolomic, and histological analyses suggested BT2 repressed a pro-ferroptotic signature in the lungs of MCT rats.

### A Low BCAA Diet Mitigated PAH and Restructured Systemic Metabolism in Monocrotaline Rats

Because BCAAs are essential amino acids^37^, we assessed how a nonpharmacological intervention using a low BCAA diet modulated PAH severity. Although not as potent as BT2, a low BCAA diet imparted notable therapeutic effects in MCT rats. In particular, the dietary intervention reduced RVSP, mPAP, and TPRi (**Figure 5A-C)**. Histological analysis also revealed diminished RV fibrosis in the MCT-low BCAA diet animals (**Figure 5D**). Next, the low BCAA diet improved RV function as TAPSE, right ventricular free wall thickness change, and cardiac output normalized to body weight were all increased (**Figure 5E-G**). Because dietary BCAAs alter metabolism in several organs, we profiled serum metabolomics to gain additional insights into how this dietary intervention impacted systemic metabolism. The low BCAA diet did not significantly decrease all BCAAs in the serum, which may have contributed to its less robust effects as compared to BT2 (**Figure 5H**). However, it did alter the systemic metabolome (**Figure 5I**) and unbiased random forest classification identified the ratio of homoarginine to symmetric dimethylarginine and predicted rate of homoarginine synthesis as important metabolites in distinguishing the MCT-Standard and MCT-Low BCAA diet rodents (**Figure 5J**).

**Figure 5:**
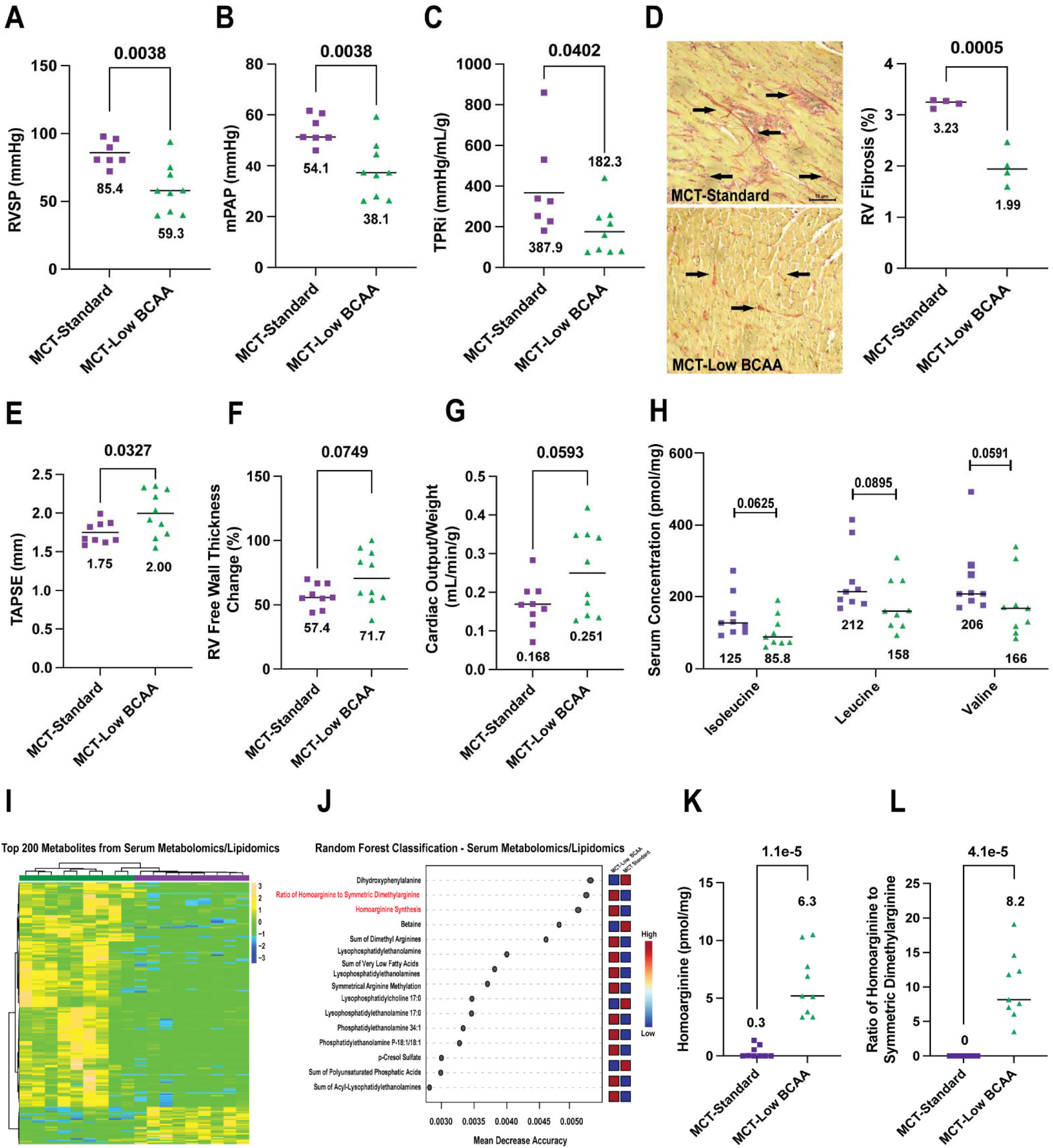
**A low BCAA diet imparted mild therapeutic effects in monocrotaline rats**. Low BCAA diet significantly reduced (A) right ventricular systolic pressure (RVSP, MCT-standard diet [MCT-Standard]: 85.4±9.4 [n=10], MCT-low BCAA diet [MCT-Low BCAA]: 59.3±18.0 mmHg [n=10]), (B) mean pulmonary artery pressure (mPAP, MCT-Standard: 54.1±5.7, MCT-Low 38.1±11.0, mmHg) and (C) total pulmonary resistance index (TPRi, MCT-Standard: 387.9±236.6, MCT-Low BCAA: 182.3±122.3 mmHg/mL/g); *p*-values determined by unpaired *t* test. These pulmonary vasculature alterations were matched by significant improvements in (D) RV fibrosis (MCT-Standard: 3.23±0.08, MCT-Low BCAA: 1.99±0.36 %, RV sections stained with Picrosirius Red) and (E) tricuspid annual plane systolic excursion (MCT-Standard: 1.75±0.14, MCT-Low BCAA: 2.00±0.29 mm), with noticeable, but not statistically significant, alterations in (F) RV free wall thickening (MCT-Standard: 57.4±9.1, MCT-Low BCAA: 71.7±20.8 %) and cardiac output normalized to body weight (MCT-Standard: 0.168±0.060, MCT-Low BCAA: 0.251±0.109 mL/min/g) (G); *p-*values determined by unpaired *t* test. (H) Serum concentrations of isoleucine (MCT-Standard: 125.0±60.8, MCT-Low BCAA: 85.8±44.3 pmol/mg) and valine (MCT-Standard: 206.0±101.0, MCT-Low BCAA: 166.0±89.6 pmol/mg) were non-significantly reduced by the Low BCAA diet, with leucine concentrations unchanged; *p*-values determined by Mann-Whitney *U* test. (I) Hierarchical cluster analysis of the top 200 serum metabolites/lipids from MCT-Standard Diet (n=9) and Low BCAA diet rodents (n=9). (J) Random forest classification of top serum metabolites and lipids important for distinguishing MCT-Standard Diet and Low BCAA diet rodents. (K) Low BCAA diet significantly increased serum homoarginine concentrations (MCT-Standard: 0.3±0.5, MCT-Low BCAA: 6.3±2.8 pmol/mg) and (L) the calculated ratio of homoarginine to symmetric dimethylarginine (MCT-Standard: 0±0, MCT-Low BCAA: 8.2±4.8); *p*-values determined by unpaired *t* test and Mann-Whitney *U* test, respectively.

Notably, the low BCAA diet increased both homoarginine levels and the ratio of homoarginine to symmetric dimethylarginine (**Figure 5K and L**). In total, these data supported the hypothesis that modulating dietary BCAAs can impact pulmonary vascular disease and indicated that a low BCAA diet may be a nonpharmacological approach to combat pulmonary vascular remodeling.

### BT2 Combatted Liver Histological, Metabolomic, and Proteomic Dysregulation in Monocrotaline Rats

Next, we investigated how BT2-mediated correction of PAH severity impacted hepatic hemodynamic stress and hepatic histological/molecular phenotypes in our three experimental groups. BT2 treatment significantly reduced maximal right atrial pressures (**Figure 6A**). Furthermore, histological evaluation of liver sections revealed an increase in hepatocyte nuclear size, a readout of shear stress^17^, in MCT rats, which was partially corrected by BT2 (**Figure 6B**). To further probe markers of shear stress, we used immunoblots to assess hepatic expression of KLF2^38^ and pAkt^39^. KLF2 expression was elevated and the ratio of pAkt to Akt were reduced in MCT-Vehicle livers, but both markers were normalized by BT2 treatment (**Figure 6C**). Likewise, BT2 partially mitigated the accumulation of shear stress response proteins in proteomics analysis (**Figure 6D**).

**Figure 6:**
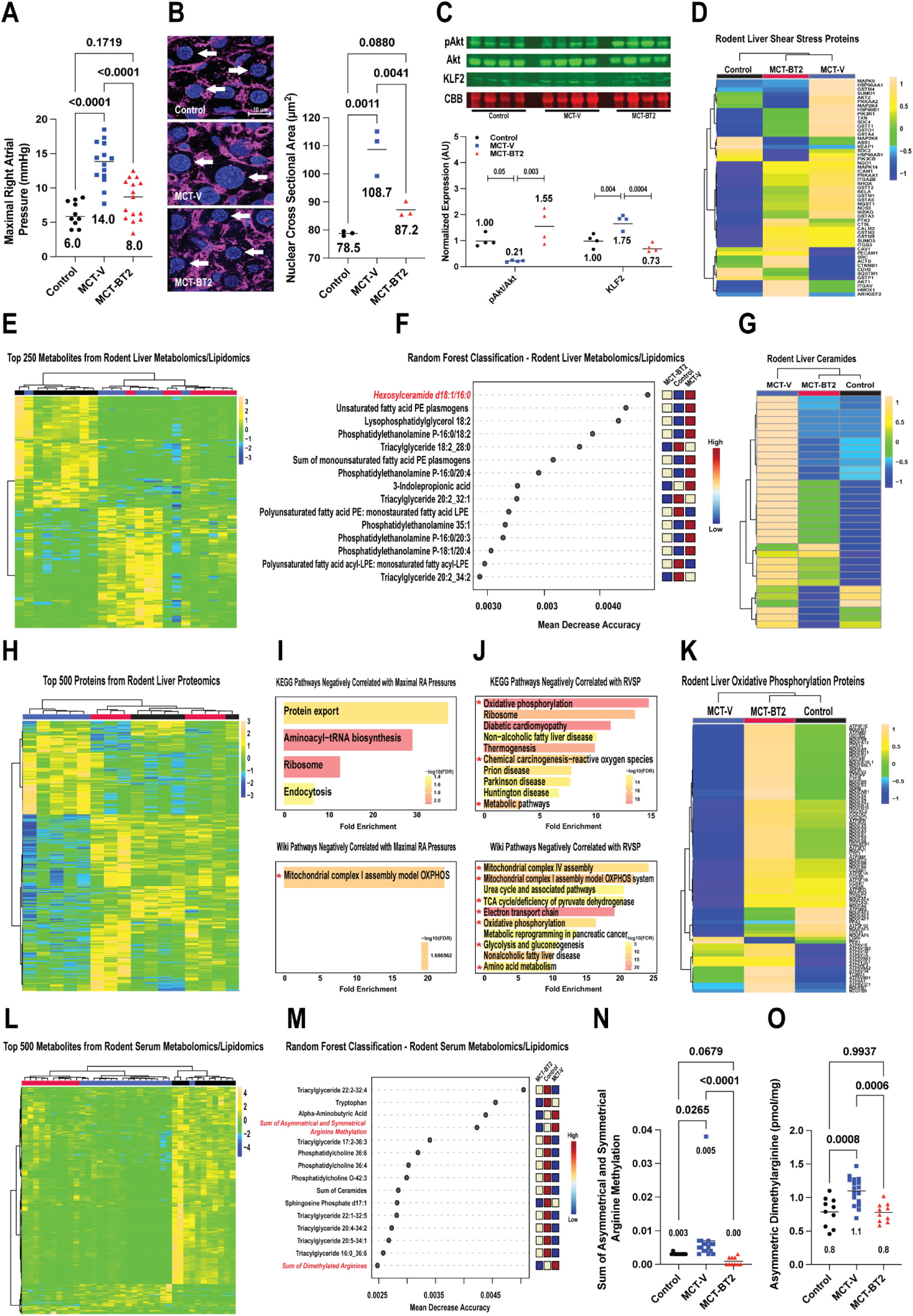
BT2 treatment mitigated activation of hepatic shear-stress phenotypes and associated metabolic derangements. (A) BT2 reduced maximal right atrial pressures (Con: 6.0±2.0, MCT-V: 14.0±3.0, MCT-BT2: 8.0±3.0 mmHg; *p-*values determined by ordinary one-way *ANOVA* with Tukey’s multiple comparison test). (B) Hepatocyte nuclear cross-sectional area was decreased in MCT-BT2 livers (Con: 78.5±0.8, MCT-V: 108.7±8.3, MCT-BT2: 87.2±2.6 μm^2^; *p*-values determined by ordinary one-way *ANOVA* with Tukey’s multiple comparison test). Nuclei are stained with WGA and DAPI. Arrows indicate hepatocyte nuclei. (C) Normalized expression of the ratio of phosphorylated Akt to Akt (Con: 1.00± 0.24, MCT-V: 0.214± 0.03, MCT-BT2: 1.55±0.65) and KLF2 (Con: 1.17±0.25, MCT-V: 1.75±0.27, MCT-BT2: 0.86±0.16) were reduced by BT2 treatment; *p-*values determined by ordinary one-way *ANOVA* with Tukey’s multiple comparison test. (D) Hierarchical cluster analysis of proteins involved in the shear stress pathway. (E) Hierarchical cluster analysis of top 250 metabolites and lipids from control (n=8), MCT-V (n=8), and MCT-BT2 (n=8) livers. (F) Random forest classification of top metabolites and lipids important for distinguishing between control, MCT-V, and MCT-BT2 livers. Hierarchical cluster analysis of (G) liver ceramides and (H) top 500 proteins from control (n=5), MCT-V (n=5), and MCT-BT2 (n=6) livers. KEGG and Wiki pathway analysis of proteins negatively correlated with (I) maximal right atrial pressure and (J) right ventricular systolic pressure. Hierarchical cluster analysis of (K) proteins in the oxidative phosphorylation pathway and (L) top 500 metabolites and lipids from control (n=10), MCT-V (n=17), and MCT-BT2 (n=10) serum samples. (M) Random forest classification of top metabolites and lipids important for differentiating between control, MCT-V, and MCT-BT2 serum. BT2 treatment significantly reduced (N) sum of asymmetric and symmetric arginine methylation (Con: 0.003±0.0003, MCT-V: 0.005±0.0081, MCT-BT2: 0.00±0.0012) and (O) asymmetric dimethylarginine (Con: 0.8±0.2, MCT-V: 1.1±0.2, MCT-BT2: 0.8±0.1 pmol/mg); *p-*values determined by Kruskal Wallis test with Dunn’s multiple comparison test and ordinary one-way *ANOVA* with Tukey’s multiple comparison test, respectively.

To determine whether these structural and protein-level changes coincided with broader liver metabolic reprogramming, we conducted multiomic profiling of rodent livers. Hierarchical clustering of the top 250 liver metabolites and lipids confirmed MCT-BT2 possessed a distinct metabolome relative to control and MCT-Vehicle livers (**Figure 6E**). Given that the liver is a physiologically important site for BCAA degradation^40^, we assessed whether hepatic BCAA metabolism was altered by BT2 treatment. Metabolomic and proteomic analyses revealed liver BCAA concentrations were not different across the three groups (**Supplemental Figure 3A**), and the abundances of BCAA degradation proteins were reduced in both MCT-Vehicle and MCT-BT2 compared to control livers (**Supplemental Figure 3B**). Notably, hexosylceramide d18:1/16:0 was identified as the most important liver metabolite for differentiating the three groups (**Figure 6F**) and overall, nearly all ceramide species abundances were elevated in MCT livers, which BT2 treatment negated (**Figure 6G**).

Because ceramides are a known regulator of mitochondrial homeostasis^41^, we then focused on mitochondria-specific alterations linked to PAH severity. Proteomic analysis of liver mitochondrial enrichments confirmed MCT-BT2 livers had a unique proteomic signature compared to both control and MCT-Vehicle livers (**Figure 6H**). Next, to better understand the liver proteomic restructuring associated with PAH severity and RV dysfunction, we performed pathway enrichment analysis on proteins negatively correlated with RVSP and maximal right atrial pressures, which identified predominantly oxidative phosphorylation/electron transport chain pathways (**Figure 6I and J**). Importantly, BT2 preserved abundances of oxidative phosphorylation proteins, in contrast to downregulation in MCT-Vehicle livers (**Figure 6K**).

Finally, to evaluate the potential systemic metabolic effects of altered hepatic function, we profiled the serum metabolome of all three groups. Hierarchical clustering of serum metabolites revealed BT2 treatment modestly altered the serum metabolome (**Figure 6L**). Random forest classification identified serum asymmetrical and symmetrical arginine methylation and the sum of dimethylated arginines as important components in differentiating the three groups (**Figure 6M**). Further analysis revealed BT2 treatment blocked the predicted sum of asymmetrical and symmetrical arginine methylation (**Figure 6N**) and normalized asymmetrical dimethylarginine levels (**Figure 6O**).

### PAH Livers Displayed Hepatocyte Nuclear Expansion and Metabolic Derangements

To examine the human relevance of our rodent hepatic phenotypes, we performed a combined histological and proteomic analysis of control (*n*=4) and PAH (*n*=3) liver samples. In agreement with our rodent data, there was a significant increase in nuclear size in PAH hepatocytes (**Figure 7A**). Next, we analyzed liver metabolic regulation by evaluating the proteome of mitochondrial enrichments of the control and PAH livers.

**Figure 7:**
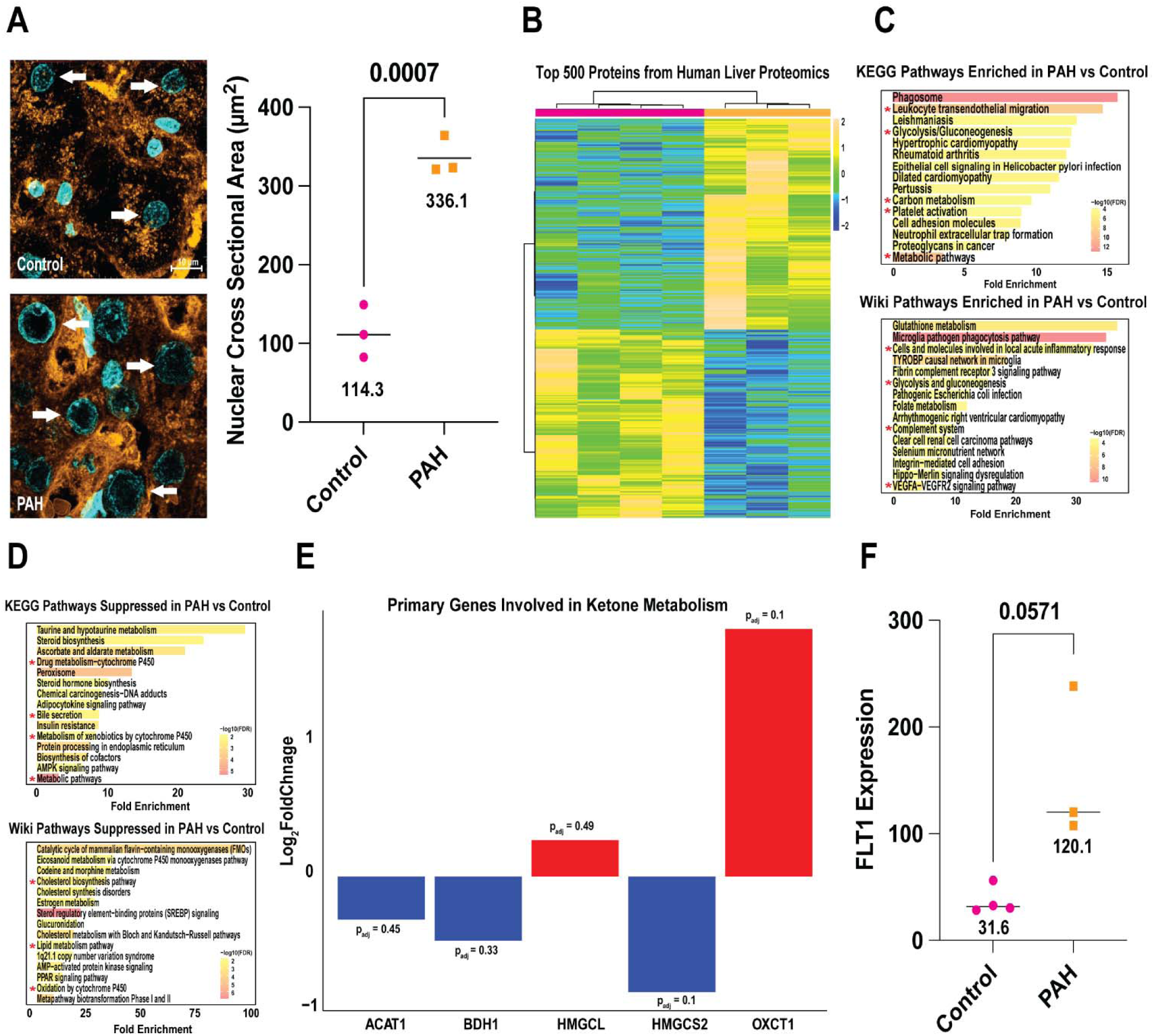
**Livers from PAH patients exhibited hepatocyte nuclear expansion and metabolic dysregulation**. (A) Confocal micrographs of liver sections stained with WGA and DAPI. PAH patients had a significant increase in mean nuclear cross-sectional area. Nuclei are stained with WGA and DAPI. Arrows indicate hepatocyte nuclei. (Control: 114.3±33.3, PAH: 336.1±24.3 μm^2^). *p-*value determined by unpaired *t* test. (B) Hierarchical cluster analysis of top 500 proteins from mitochondrial enrichments of control (n=4) and PAH (n=3) livers. (C) KEGG and Wiki pathway analysis of proteins upregulated in PAH samples as determined by a false-discovery rate of 0.1. (D) KEGG and Wiki pathway analysis of proteins downregulation in PAH samples determined by a false-discovery rate of 0.1. (E) Key genes involved in ketone metabolism were altered in PAH livers. (F) FLT1 expression was non-significantly increased in PAH livers (Control: 31.6±12.9, PAH: 120.1±72.2; *p-*value determined by Mann-Whitney *U* test).

Hierarchical clustering of the top 500 liver proteins demonstrated the proteomic signature of PAH was distinct from controls (**Figure 7B**). Pathway analysis of significantly enriched proteins in PAH (FDR: 0.1 due to small sample size) revealed upregulation of pro-inflammatory pathways, glycolysis, and vascular endothelial growth factor (VEGF) signaling (**Figure 7C**). In contrast, pathway analysis of suppressed proteins indicated significant dysregulation of the liver’s metabolic capabilities, with reductions in cytochrome P450, bile secretion, and lipid metabolism pathways (**Figure 7D**). In accordance with the liver’s crucial role in ketogenesis, we then evaluated the abundances of several proteins involved in ketone metabolism^42,43^. Five ketogenic proteins were identified in our liver proteomics dataset, with three downregulated (ACAT1, BDH1, and HMGCS2) and two upregulated (HMGCL, OXCT1) in PAH livers, although these changes failed to reach statistical significance (**Figure 7E**).

Besides ketogenesis, the liver can also impact systemic metabolism through the production of vasoactive mediators. One notable protein associated with pulmonary vascular remodeling^44^ is Fms related receptor tyrosine kinase 1 (FLT1, a receptor for the pro-angiogenic VEGF pathway^45^), and FLT1 was elevated in PAH livers (**Figure 7F**). Collectively, these data demonstrated PAH livers displayed altered nuclear morphology, dysregulation of numerous metabolic pathways, and heightened levels of FLT1.

## Discussion

Here, we provide multiple lines of evidence demonstrating disruptions in BCAA metabolism impact PASMC biology and pulmonary vascular disease. Starting with human data, we confirm the presence of altered BCAA metabolites across different physiological gradients in a cohort of PAH and chronic thromboembolic pulmonary hypertension patients. Then, in lung transcriptomic studies, we show dysregulation of the BCAA catabolic and ferroptosis pathways in human PAH PASMCs. *In vitro*, excess BCAAs induce pro-ferroptotic phenotypes in non-diseased human PASMC. Next, preclinical rodent studies show that augmenting BCAA catabolism dampens PAH severity, improves RV function, and enhances exercise capacity. Additionally, we demonstrate BT2 treatment mitigates lung BCAA accumulation, rescues lung-specific metabolic deficits in BCAA metabolism and combats mTORC1 activation. Multi-omic and targeted histological analyses suggest that altered ferroptosis drives these therapeutic benefits as BT2 normalizes several metabolites with ties to ferroptosis and blunts perivascular complement deposition and transferrin receptor accumulation in PASMCs. We then show a nonpharmacological approach imparts beneficial effects as a low BCAA diet lessens disease severity and alters systemic metabolism in MCT rats. Furthermore, in MCT livers, we demonstrate that administering BT2 reduces shear stress signaling and associated changes in hepatocyte nuclear morphology, mitigates mitochondrial dysfunction, and blunts modifications to the systemic metabolome. Importantly, we also confirm that human PAH livers recapitulate the hepatocyte nuclear expansion phenotype observed in MCT livers and identify several dysregulated metabolic pathways, including suppression of the cytochrome P450 system and altered ketone metabolism. In conclusion, these data implicate impaired PASMC BCAA metabolism as an important component of lung PAH pathogenesis via ferroptosis and suggest PAH severity impacts liver metabolism through hepatic mechanical stress phenotypes.

Our manuscript indicates that targeting disruptions in BCAA catabolism and ferroptosis induction may have therapeutic relevance, and this hypothesis could be addressed with a currently available pharmaceutical. 4-phenylbutyrate (4PBA), an ammonia scavenger commonly prescribed for urea cycle disorders^46^, is an effective inducer of BCAA catabolism. In acute lung injury, 4PBA treatment reduces pulmonary inflammation and ferroptosis^47^. These results align with our data, in which inducing BCAA catabolism via BT2 blunts classical complement and ferroptosis activation in the lungs. Interestingly, 4PBA treatment mitigates PAH severity in both the chronic hypoxia^48^ and monocrotaline^49^ models of PAH. While these data highlight the potential utility of activating BCAA catabolism in PAH, longitudinal clinical studies are still needed. Currently, there is an ongoing phase II clinical trial evaluating 4PBA in idiopathic or hereditary PAH^50^ and hopefully this trial will provide further insights regarding the role of BCAA metabolism in PAH pathobiology.

These data add to a growing body of literature showing BCAA metabolism plays a key role in vascular biology. BCAA catabolic deficits are observed in thoracic aortic dissection (TAD), resulting in vascular smooth muscle cell phenotype switching via mTOR overactivation^51^. Consistent with our results, BT2 treatment in TAD animals reactivates BCAA catabolism and reduces mTOR activation, ultimately blocking pathological vascular remodeling. In addition, impairments in BCAA oxidation are linked to systemic hypertension. Murashige *et. al.* shows that BT2 administration in post myocardial infarction rodents is cardioprotective, with reductions in blood pressure and improvements in vascular relaxation^52^. In support of these findings, a Mendelian randomization study demonstrates that elevated plasma BCAAs predict higher blood pressure in humans. A related clinical trial reports similar results, as increased circulating BCAAs at the study entry strongly correlate with subsequent incident hypertension^53^. Our findings alongside these other manuscripts demonstrate a crucial role of BCAA metabolism in vascular homeostasis.

Our data further position ferroptosis as a key regulator of pulmonary vascular disease. We showed ferrostatin-1 reduces PAH severity, mitigates ectopic complement deposition, and exerts potent anti-inflammatory effects in the pulmonary vasculature^6^. These findings reinforce several independent studies that link ferroptosis to PAH across diverse experimental models. *In vitro*, Lin *et. al.* demonstrates ca-circSCN8A, a pro-ferroptotic circular RNA, is upregulated in pulmonary hypertension, and its overexpression induces ferroptosis in human PASMCs^54^. Similarly, another study finds that a different circular RNA, circMyst4, decreases under hypoxia and that restoring circMyst4 alleviates PASMC ferroptosis^55^. Finally, large-scale bioinformatic analyses of human pulmonary vascular tissues reveal altered expression of multiple ferroptosis-associated genes in PAH^56,57^. Thus, these data highlight the role that ferroptosis has in pathological remodeling in the pulmonary vasculature.

This manuscript adds to an emerging dataset showing multiple amino acids regulate pulmonary vascular disease. Xiao *et. al*. show BCKAs activate HIF-1 in PASMCs, and these data are matched with elevated systemic levels of BCKAs in PAH patients^2^. In addition, glycine drives PAH pathogenesis via BOLA3 deficiency, resulting in endothelial cell proliferation and vasoconstriction in the pulmonary vasculature^58^.

Moreover, glutamine and serine worsen disease progression in PAH by promoting collagen biosynthesis, which fuels pulmonary fibroblasts and vascular fibrosis^59^. Proline metabolism is critical for supporting biomass generation during small vessel pulmonary arterial remodeling in PAH^60^. In the Sugen/hypoxia model, inhibiting tryptophan metabolism blunts pulmonary artery hypermuscularization and reduces cardiac fibrosis^61^. Together, our findings and others highlight the key role of amino acid metabolism in pulmonary vascular remodeling.

Next, our work better characterizes an emerging and biologically important lung-right heart-liver axis in PAH. The liver is an increasingly relevant component of PAH pathobiology, as even minor changes in liver functional assessments are highly prognostic^14^ and PAH patients display elevated liver stiffness^15^, which strongly correlates with RV failure. One potential trigger for PAH-hepatology is hemodynamic shear stress as tricuspid regurgitation propels retrograde blood flow into the right atrium and inferior vena cava^62^, which could expose downstream organs to turbulent venous flow, disrupt the phospholipid bilayer^63^, and induce sphingolipid production^64^. Sphingomyelinases cleave sphingolipids into ceramides, which then accumulate in the mitochondria and destabilize the electron transport chain^65–67^, potentially accounting for the reduced oxidative phosphorylation in MCT-Vehicle livers. Then, we nominate a potential biomarker for this process as serum asymmetric dimethylarginine (ADMA) is elevated in MCT-Vehicle animals. The liver primarily metabolizes ADMA and high circulating levels are found in liver cirrhosis, alcoholic hepatitis, and acute liver failure^68^. Thus, our data reinforce the presence of liver metabolic dysfunction in PAH. However, future comparisons of liver molecular phenotypes across different disease states may further clarify how PAH specifically impacts the liver.

Given the liver’s central metabolic role in systemic biology, the observed hepatic alterations may have broader consequences for PAH. The liver is the sole producer of ketones, an important systemic fuel source with potent anti-inflammatory effects^43^. In a prior study, we show PAH patients with decompensated RV failure lack a compensatory ketosis^69^. Here, we demonstrate that ketogenic proteins are altered in PAH livers, providing additional evidence for disrupted ketone metabolism. In addition, PAH livers show elevated expression of FLT1, a receptor for the pro-angiogenic VEGF pathway^45^. FLT1 is a biomarker in PAH, with serum levels strongly correlating with disease severity^44^. Liver-derived FLT-1 may contribute to high circulating levels in PAH and ultimately promote pulmonary vascular remodeling. Collectively, these findings position the liver as an active contributor to PAH pathogenesis, with effects that extend beyond local dysfunction to shape systemic and pulmonary vascular outcomes.

## Limitations

This study has several important limitations that we must acknowledge. First, we only evaluated male rodents due to a more robust PAH phenotype. Then, a previous study showed BCAAs were decreased across the pulmonary gradient in the Sugen/hypoxia PAH model^70^. However, this discrepancy could be due to differences in disease severity induced by the respective models, as Lin et. al. reported serum leucine and isoleucine were increased in MCT rodents^71^. Next, we used whole lung tissue for our proteomic and metabolomic analyses and thus it is possible that altered BCAA metabolism and ferroptosis could be occurring in cell types other than PASMC. We tried to overcome some of these limitations by including the human lung transcriptomic data, which showed that SMC abundance had a strong negative correlation with ferroptosis pathway activity and PAH PASMC had significant deficits in the BCAA catabolic and ferroptosis pathways relative to healthy controls. Furthermore, because human PAH tissue is rare and difficult to obtain, there was a small sample size (control: n=4, PAH: n=3) for the liver proteomics analysis, thus the FDR for pathway enrichment was increased to 0.1 to identify additional pathways that may be of interest for future investigations. Finally, it is possible that shear stress is not the only driver of liver dysfunction as PAH compromises cardiac output^72^ and is associated with systemic metabolic derangements^73^.

## Supplemental Materials

Figures S1-S3 Supplemental Table 1 Data Sets

## Non-Standard Abbreviations and Acronyms

4PBA: 4-phenylbutyrate
ART: Arterial
ADMA: Asymmetric dimethylarginine
BCAA: Branched-chain amino acids
BCAT2: Branched-chain amino acid aminotransferase 2
BT2: 3,6-Dichloro-1-benzothiophene-2-carboxylic acid
DAPI: 4’,6-diamidino-2-phenylindole
FDR: False discovery rate
HIF1: Hypoxia-inducible factor 1
MCT: Monocrotaline
mPAP: Mean pulmonary arterial pressure
MS: Mass spectrometry
mTOR: Mammalian target of rapamycin
mTORC1: Mammalian target of rapamycin complex 1
PA: Pulmonary artery
PAH: Pulmonary arterial hypertension
PASMC: Pulmonary artery smooth muscle cells
RNAseq: RNA-sequencing
ROS: Reactive oxygen species
RV: Right ventricular/right ventricle
RVSP: Right ventricular systolic pressure
SMC: Smooth muscle cell
snRNAseq: Single nucleus RNA-sequencing
SVC: Superior vena cava
TAPSE: Tricuspid annular plane systolic excursion
TAD: Thoracic aortic dissection
TPRi: Total pulmonary resistance index
VEGF: Vascular endothelial growth factor
WGA: Wheat germ agglutinin

---

**Supplemental Figure 1:**
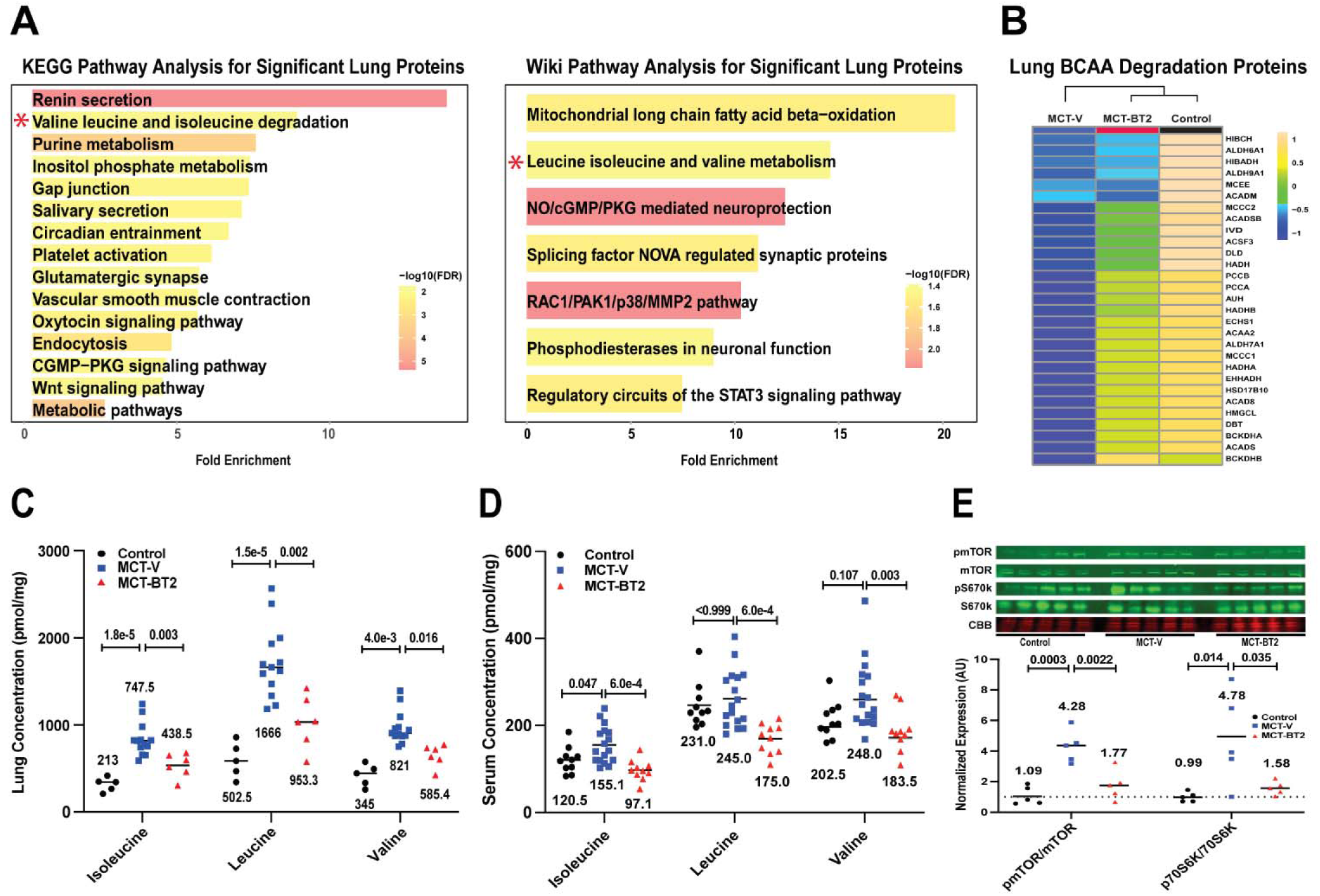
BT2 treatment blunted metabolic dysregulation and mTOR activation in the lungs. (A) KEGG and Wiki pathway analysis of statistically significant lung proteins from mitochondrial enrichments of control (n=5), MCT-V (n=13), and MCT-BT2 lung tissue (n=6). *p-*values determined by ordinary one-way *ANOVA* with Tukey’s multiple comparison test. (B) Hierarchical cluster analysis of proteins in the KEGG isoleucine, leucine, and valine degradation pathway. Red text denotes statistically significant proteins, with *p-*values determined by ordinary one-way *ANOVA* with Tukey’s multiple comparison test. (C) BT2 treatment significantly reduced lung concentrations of isoleucine (Con: 213.0±83.5, MCT-V: 747.5± 196.0, MCT-BT2: 438.5±143.1 pmol/mg), leucine (Con: 502.5±211.0, MCT-V: 1666.0±429.7, MCT-BT2: 953.3±313.6 pmol/mg), and valine (Con: 345.0±130.4, MCT-V: 821.0±194.2, MCT-BT2: 585.4±131.2 pmol/mg). *p-*values determined by ordinary one-way *ANOVA* with Tukey’s multiple comparison test for isoleucine and leucine, Kruskal Wallis test with Dunn’s multiple comparison test for valine. (D) BT2 significantly reduced isoleucine (Con: 120.5±30.4, MCT-V: 155.1±42.3, MCT-BT2: 97.1±23.9 pmol/mg), leucine (Con: 231.0±51.3, MCT-V: 245.0±65.4, MCT-BT2: 175.0±34.6 pmol/mg), and valine (Con: 202.5±42.1, MCT-V: 248.0±77.9, MCT-BT2: 183.5±48.3 pmol/mg) serum concentrations, but only isoleucine was significantly different between control and MCT-V. *p-*values determined by Kruskal Wallis test and Dunn’s multiple comparison test for valine and leucine, and ordinary one-way *ANOVA* with Tukey’s multiple comparison test for isoleucine. (E) Normalized expression of the ratio of phosphorylated mTOR to mTOR (Con: 1.09±0.59, MCT-V: 4.28±1.07, MCT-BT2: 1.77±0.97) and phosphorylated 70S6K to 70S6K (Con: 0.99±0.31, MCT-V: 4.78±3.0, MCT-BT2: 1.58±0.44). Induction of BCAA catabolism significantly reduced pulmonary mTOR activation. *p-*values determined by ordinary one-way *ANOVA* with Tukey’s multiple comparison test.

**Supplemental Figure 2:**
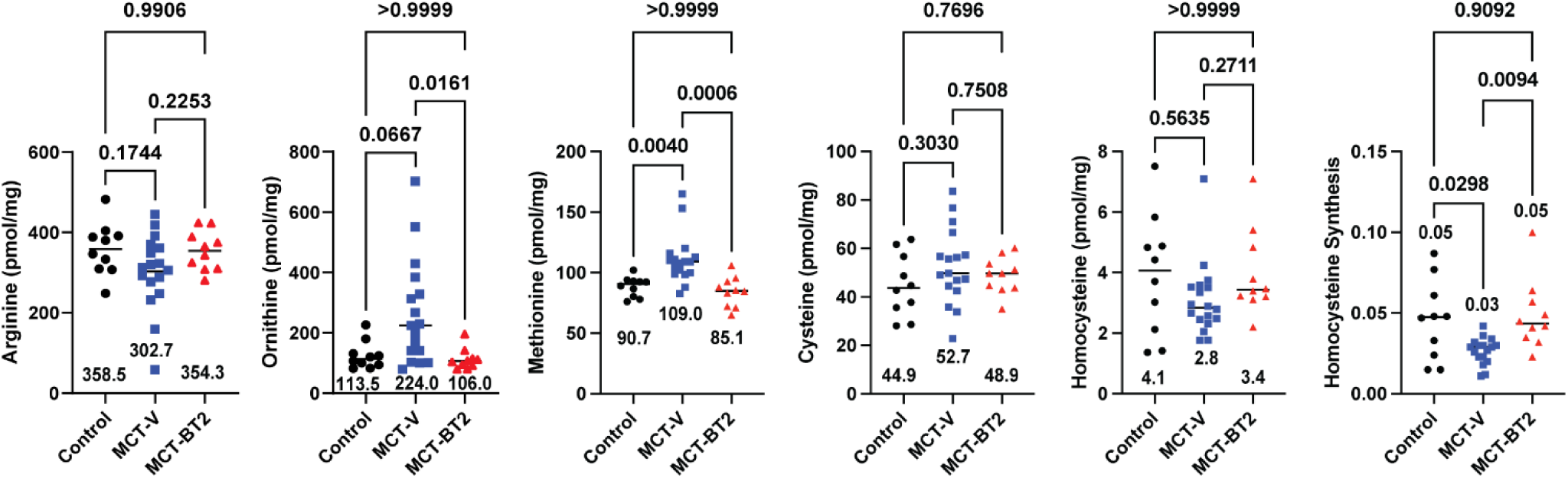
Serum metabolites do not fully recapitulate therapeutic alterations observed in BT2 lungs. Ornithine (Con: 113.5±45.6, MCT-V: 224.0±173.6, MCT-BT2: 106.0±34.4) and methionine (Con: 90.7±8.2, MCT-V: 109.0±20.4, MCT-BT2: 85.1±12.4) were significantly reduced in MCT-BT2 serum relative to MCT-V, while the predicted rate of homocysteine synthesis (Con: 0.05±0.02, MCT-V: 0.03±0.01, MCT-BT2: 0.05 ±0.02) was suppressed in MCT-V. Serum concentrations of arginine (Con: 358.8±64.8, MCT-V: 302.7±94.0, MCT-BT2: 354.3±49.2), cysteine (Con: 44.9±12.8, MCT-V: 52.7±15.6, MCT-BT2: 48.9±7.6), and homocysteine (Con: 4.1±2.0, MCT-V: 2.8±1.2, MCT-BT2: 3.4±1.4) were unchanged. *p-*values determined by: Kruskal Wallis test with Dunn’s multiple comparison test for ornithine, methionine, and homocysteine; ordinary one-way *ANOVA* with Tukey’s multiple comparison test for arginine, cysteine, and homocysteine synthesis.

**Supplemental Figure 3:**
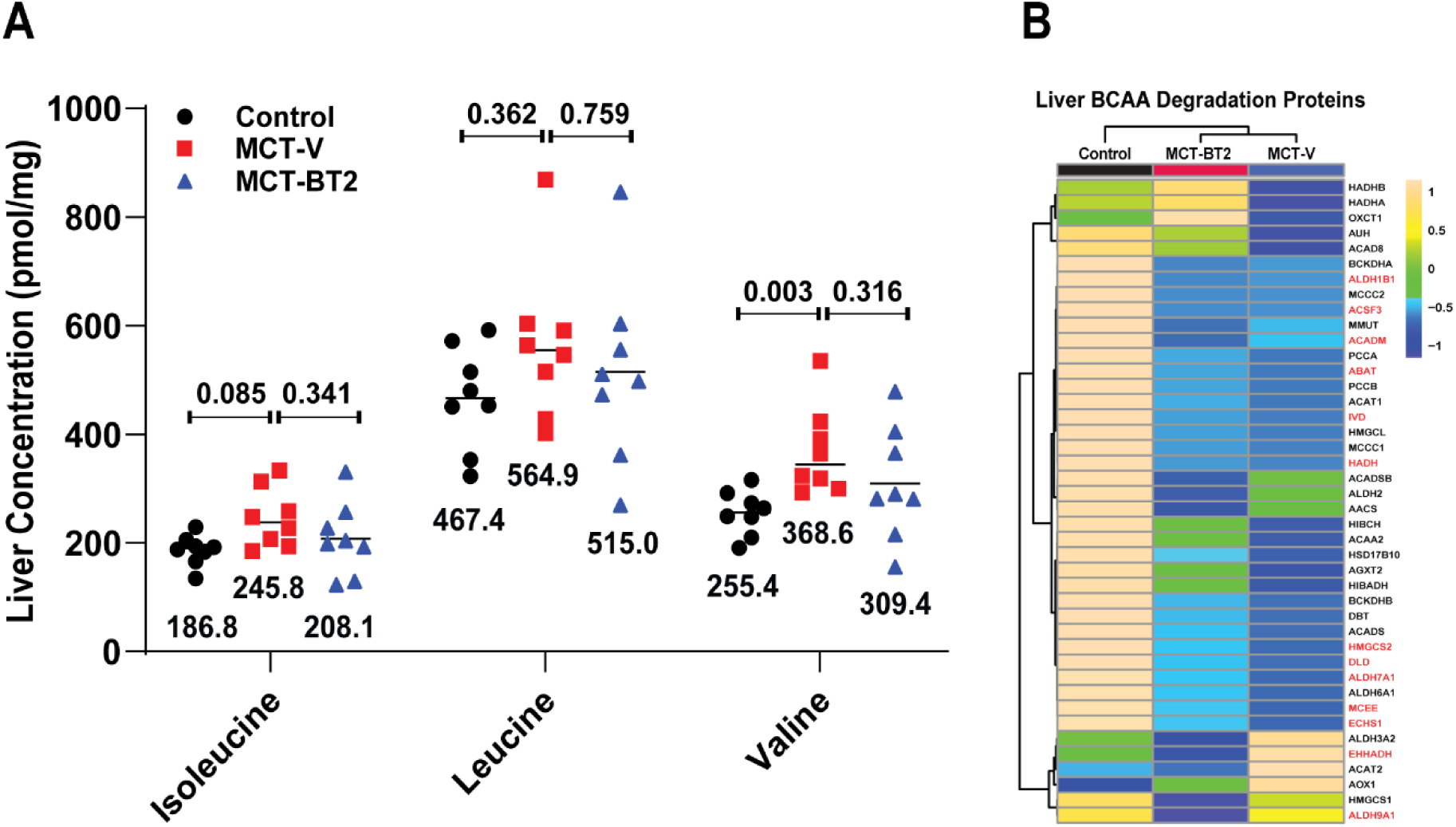
Hepatic BCAA metabolism was only modestly altered by BT2 treatment. (A) Isoleucine (Con: 186.8±27.7, MCT-V: 245.8±54.3, MCT-BT2: 208.1±67.2 pmol/mg) and leucine (Con: 467.4±95.1, MCT-V: 564.9±142.8, MCT-BT2: 515.0±171.0 pmol/mg) concentrations in the liver were not different among control, MCT-V, and MCT-BT2, while valine (Con: 255.4±40.9, MCT-V: 368.6±81.1, MCT-BT2: 309.4±103.7 pmol/mg) was only significantly different between control and MCT-V liver samples. *p-*values determined by ordinary one-way *ANOVA* with Tukey’s multiple comparison. (B) Hierarchical cluster analysis of proteins in the isoleucine, leucine, and valine degradation pathway. Red text denotes statistically significant proteins, with *p-*values determined by ordinary one-way *ANOVA* with Tukey’s multiple comparison test.

**Supplemental Table 1.** Antibodies Used for Immunoblots and Confocal Microscopy.

| Antigen | Company | Species | Dilution |
| --- | --- | --- | --- |
| **Western Blots** |  |  |  |
| *Lung* |  |  |  |
| mTOR | Cell Signaling | Rabbit | 1:250 |
| Phospho-mTOR | Cell Signaling | Rabbit | 1:250 |
| 70S6K | Cell Signaling | Rabbit | 1:250 |
| Phospho-70S6K | Cell Signaling | Rabbit | 1:250 |
| *Liver* |  |  |  |
| KLF2 | Abcam | Rabbit | 1:250 |
| Akt | Cell Signaling | Rabbit | 1:250 |
| Phospho-Akt | Cell Signaling | Rabbit | 1:250 |
| Anti-rabbit 800 secondary | Li-COR | Goat | 1:5000 |
| **Immunofluorescence** |  |  |  |
| C3 | Abcam | Rabbit | 1:50 |
| Transferrin receptor | Abcam | Rabbit | 1:50 |
| FITC conjugated $\alpha_{sm}$-actin | Sigma Aldrich | Mouse | 1:50 |
| Anti-rabbit Alexa Fluor<br>568 secondary | Invitrogen | Goat | 1:500 |

